# Eukaryotic genomic data uncover an extensive host range of mirusviruses

**DOI:** 10.1101/2024.01.18.576163

**Authors:** Hongda Zhao, Lingjie Meng, Hiroyuki Hikida, Hiroyuki Ogata

## Abstract

A recent marine metagenomic study has revealed the existence of a novel group of viruses designated mirusviruses, which are proposed to form an evolutionary link between two realms of double-stranded DNA viruses, *Varidnaviria* and *Duplodnaviria*. Metagenomic data suggest that mirusviruses infect microeukaryotes in the photic layer of the ocean, but their host range remains largely unknown. In this study, we investigated the presence of mirusvirus marker genes in publicly available 1,901 eukaryotic genome assemblies, mainly derived from unicellular eukaryotes, to identify potential hosts of mirusviruses. Mirusvirus marker sequences were identified in 1,348 assemblies spanning 284 genera across eight supergroups of eukaryotes. The habitats of the putative mirusvirus hosts included not only marine but also other diverse environments. Among the major capsid protein (MCP) signals in the genome assemblies, we identified 85 sequences that showed high sequence and structural similarities to reference mirusvirus MCPs. A phylogenetic analysis of these sequences revealed their distant evolutionary relationships with the seven previously reported mirusvirus clades. Most of the scaffolds with these MCP sequences encoded multiple mirusvirus homologs, underscoring the impact of mirusviral infection on the evolution of the host genome. We also identified three circular mirusviral genomes within the genomic data of the oil producing thraustochytrid *Schizochytrium* sp. and the endolithic green alga *Ostreobium quekettii*. Overall, mirusviruses probably infect a wide spectrum of eukaryotes and are more diverse than previously reported.

**Highlights:**

- Mirusvirus signals detected in genomic data from eight eukaryotic supergroups.
- Habits of putative mirusvirus hosts not limited to marine environments.
- Major capsid sequences from these assemblies show new mirusviral lineages.
- Three circular mirusvirus genomes were identified.

## Introduction

Viruses pervade diverse environments across the Earth and play crucial ecological and evolutionary roles.^1–3^ A recent metagenomic analysis identified a previously unrecognized but diverse group of double-stranded (ds) DNA viruses, designated mirusviruses, that are abundant, widespread, and active in the global marine ecosystem.^4^ Based on their virion morphogenesis genes (e.g., HK97-fold MCP), mirusviruses are proposed to form the phylum ‘*Mirusviricota*’ in the realm *Duplodnaviria*, which includes herpesviruses and caudoviruses (i.e., tailed viruses of prokaryotes). However, a large number of genes encoded by mirusviruses, including informational genes (e.g., DNA polymerase, RNA polymerase), are more closely related to homologs in nucleocytoviruses. Nucleocytoviruses belong to another dsDNA viral realm, *Varidnaviria*, and are known to play important roles in marine ecosystems. The genomic similarities between mirusviruses and nucleocytoviruses suggest an evolutionary interplay between the two distinct groups of viruses and their similar habitats.

Viral groups have different host ranges. Nucleocytoviruses infect diverse eukaryotes, including protists, green algae, and animals.^5^ Within *Duplodnavira*, caudoviruses infect a broad spectrum of prokaryotes,^6,7^ whereas herpesviruses have a relatively narrow host range, limited to animals.^8,9^ Until the discovery of mirusviruses, no duplodnaviruses were known to infect basal eukaryotes, such as protists. Mirusviruses were the first group of duplodnaviruses suggested to infect unicellular eukaryotes. This suggestion was based on the horizontal gene transfer of heliorhodopsin genes between mirusviruses and green algae, and the fact that mirusvirus signals were detected in the metagenomes and metatranscriptomes derived from size fractions corresponding to unicellular planktonic eukaryotes.^4^ Therefore, the discovery of mirusviruses was considered to fill the host gap in the duplodnaviruses (between prokaryotes and animals). However, the evidence of mirusvirus hosts was limited and the taxonomic breadth of their hosts remained poorly investigated.

Two recent studies investigating the viral signals in eukaryotic genomes revealed some potential hosts of mirusviruses. Specifically, endogenized mirusviral genomes have been detected in the thraustochytrids *Aurantiochytrium limacinum* and *Hondaea fermentalgiania* (i.e., single-celled saprotrophic eukaryotes),^10^ and the phago-mixotrophic green alga *Cymbomonas tetramitiformis.*^11^ Notably, an additional circular mirusvirus-like genome (probably in the form of an episome) was identified in the *A. limacinum* genome assembly, suggesting an episomal form of mirusvirus in the host cells.^10^ In general, the viral sequences within eukaryotic genome assemblies can be categorized into three types: transferred genes, integrated viral genome fragments, and free viral genomes.^12–14^ The previously reported endogenized mirusviral sequences and the circular genome correspond to the latter two types, respectively. These three types of evidence strongly suggest that eukaryotes containing viral signals have acted as the hosts of viruses.

In the present study, we systematically screened 1,901 genome assemblies of eukaryotes for mirusvirus signals. The genomic data cover a diverse group of predominantly cultured unicellular eukaryotes. Of the 318 eukaryotic genera analyzed, 284 contained mirusvirus signals. The mirusviral MCP sequences identified in this study formed distinct phylogenetic clades, separate from previously reported mirusviral MCP sequences derived from marine environments. Moreover, three circular mirusviral genomes encoding a nearly complete set of marker genes were identified. This study suggests that the host range of mirusviruses is broad, and that mirusviruses display previously unrecognized diversity.

## Results

### Mirusviral marker sequences detected in a wide range of eukaryotes

The dataset for this study was collected from GenBank and comprised 1,901 eukaryotic genomic assemblies. These assemblies spanned over 318 minor lineages (mostly at the genus level, and are hereafter referred to as ‘genera’ for simplicity) and 16 major lineages of eukaryotic organisms (mostly at the phylum level or higher; Supplementary Table S1, Data S1), and cover eight of the nine eukaryotic supergroups (the single exception is Hemimastigophora).^15^ In these assemblies, we identified nearly 118 million open reading frames (ORFs). Of these protein sequences, 6,659 showed significant sequence similarities (E-value < 10^−5^) to the hidden Markov models (HMMs) of five selected mirusviral marker sequences: MCP, capsid triplex subunit 1 (Triplex1), capsid triplex subunit 2 (Triplex2), capsid portal protein (Portal), and capsid maturation protease (Maturation). To ensure their specificity to the mirusviruses, we aligned these sequences with other duplodnavirus (caudovirus and herpesvirus) sequences and HMMs in the PFAM database, and excluded possible false positives. In this way, we identified 6,042 marker sequences (541 MCP, 1,202 Portal, 509 Triplex1, 625 Triplex2, and 3,165 Maturation) that are specifically related to mirusvirus counterparts in 1,348 eukaryotic genome assemblies (71% of the analyzed eukaryotes).

The 1,348 eukaryotic genome assemblies containing viral marker sequences included 284 genera spanning 15 major lineages and eight eukaryotic supergroups (Data S2). Specifically, MCP was detected in 98 genera, Portal in 171 genera, Triplex1 in 114 genera, Triplex2 in 135 genera, and Maturation in 249 genera. Of these 284 genera, we selected 90 genera that showed strong signals for mirusviral marker sequences based on the criterion that the assemblies in the genus contained four or five distinct marker sequences (Fig. 1). Of the 6,042 marker sequences identified, 4,365 belonged to these 90 genera. These genera were spread across 11 major lineages and seven supergroups of eukaryotes.

**Fig. 1.**
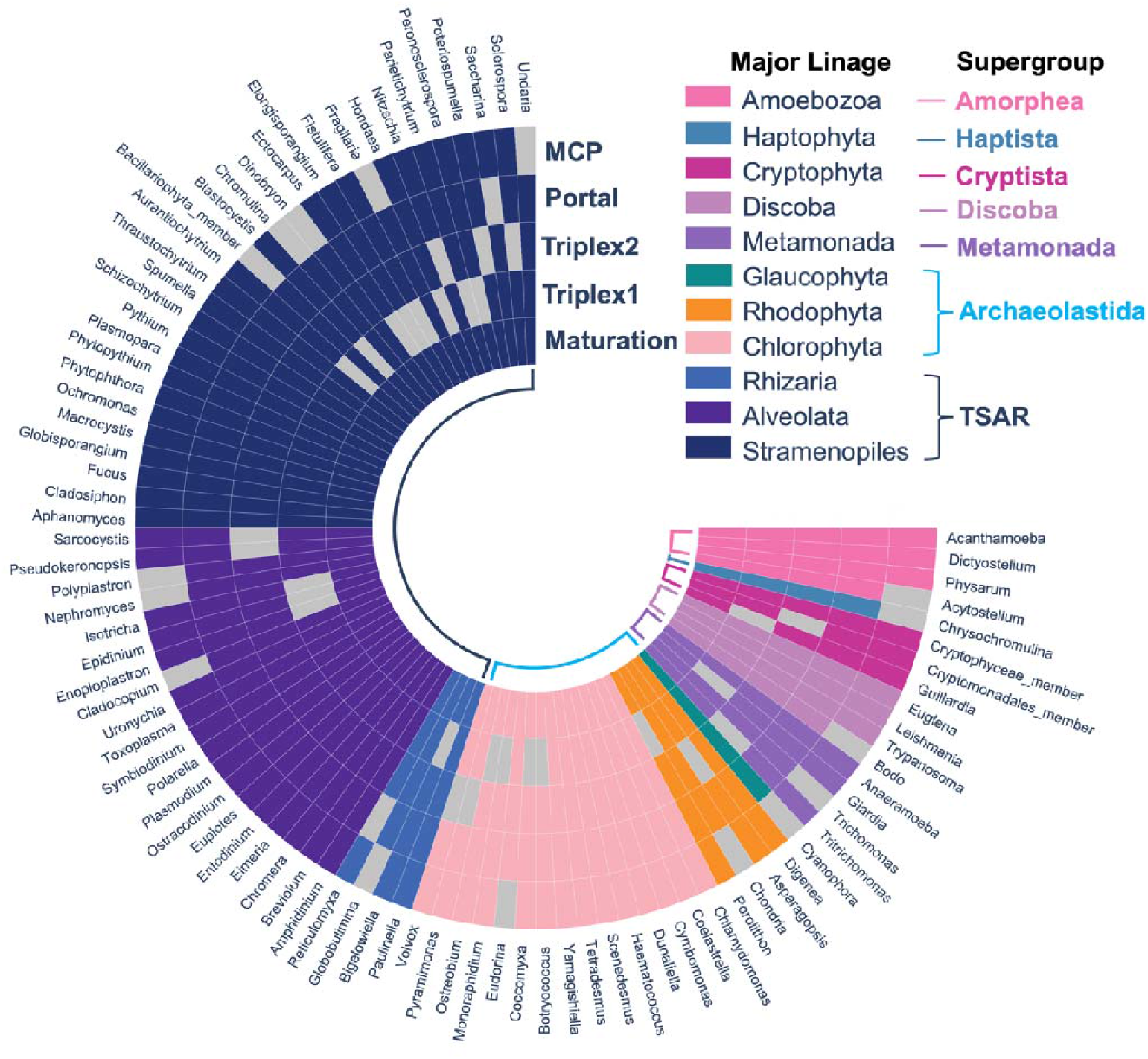
Eukaryotic genera showing four or five different mirusvirus markers in their assemblies. Colors of the cells represent the major lineages of the genus. Layers of cells represent the presence (colored) or absence (gray) of MCP, Portal, Triplex2, Triplex1, or Maturation sequences within each genus from the outermost to the innermost, respectively. Supergroups of eukaryotes are indicated in colored bold letters. Of the eight supergroups with mirusvirus signals, the supergroup CRuMs is not shown in this figure as it contained no genus with four or five mirusviral markers.

We investigated the sequence similarities between all 6,042 mirusviral marker sequences detected in the eukaryotic assemblies and reference marker sequences encoded in previously described mirusvirus genomes derived from marine metagenomes.^4^ Most of the marker sequences shared low sequence similarity scores with the reference markers, and the median bit-score calculated with hmmscan ranged from 10 to 15 for all five marker genes (Fig. 2A). The median lengths of the marker sequences in the eukaryotic assemblies were smaller than those of the reference markers and ranged from 100 to 200 amino acids (Fig. 2B). These features suggest that many of the marker sequences detected were decaying nonfunctional genes (Fig. 2C). Nevertheless, there were also marker sequences of normal length but that shared low bit scores with reference sequences, implying the existence of diversified viral lineages.

**Fig. 2.**
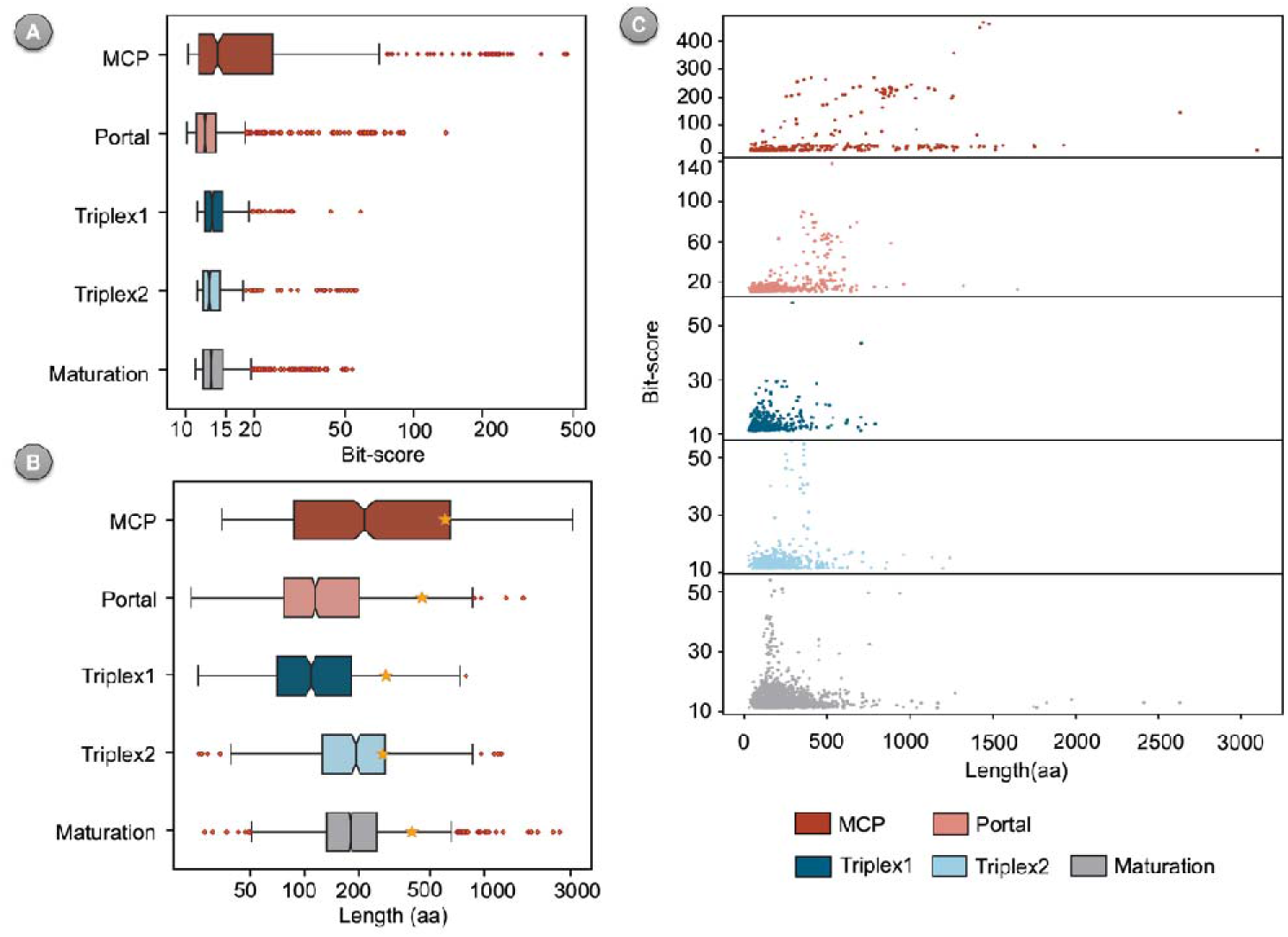
Lengths of detected marker sequences and their similarities to reference sequences. **(A)** Boxplots of bit-scores against five reference marker genes. **(B)** Boxplots of the length distributions of the detected marker sequences. Yellow stars represent the median lengths of the reference mirusvirus orthogroups. **(A–B)** Red diamonds represent the outlier points. **(C)** Relationships between length and bit-score for each marker.

### MCPs from eukaryotic assemblies form clades distinct from those of known mirusviruses

We next investigated the evolutionary relationships between the mirusvirus signals in the eukaryotic genome assemblies and previously described mirusviruses in marine metagenomes, focusing on MCP. This protein is the most important viral structural component and is critical for viral taxonomic classification.^1^ Furthermore, the tree based on MCP sequences are reported to represent the vertical evolution of mirusviruses, which is consistent with that based on informational genes.^4^ To minimize the interference degraded genes impose on phylogenetic analyses, we specifically filtered MCP marker sequences suitable for evolutionary analyses as follows. First, from the initial 541 MCP sequences identified in eukaryotic assemblies, we selected 262 sequences suitable for structural prediction. The selection was based on the absence of ambiguous amino acids (due to sequencing quality) and a length of 200–1,500 amino acids assumed to be normal for a functional MCP. We predicted the three-dimensional (3D) structures of these 262 MCPs and then compared their structural similarities with the marine mirusviral MCPs. Of the 262 structural models, 170 showed a template modeling score (TM-score) of > 0.5 (a criterion for the potential same fold^16,17^) with at least one reference mirusviral MCP, supporting their structural similarities.

We further refined our selection of MCPs from eukaryotic assemblies for phylogenetic analysis based on the conservation of a key structural feature. The HK97-fold is an important shared feature of duplodnaviral MCP structures. There are multiple conserved elements in the HK97-fold, including the axial domain (A-domain), peripheral domain (P-domain), extended loop (E-loop), and N-terminal arm (N-arm).^18^ The floor domain, which includes the last three elements, exists in all HK97 MCPs. We used the E-loop, a long two-stranded β-sheet hairpin, as an indicator of the presence of the floor domain as it is readily recognized in a predicted structure if present. Of the 170 sequences with high structural similarities to reference mirusviral MCPs, 150 were found to contain the two-stranded β-sheet hairpin with a β-strand longer than 10 amino acids on each side.

Finally, we compared the structures of these 150 MCP sequences with those of other representative duplodnaviral MCPs (from caudoviruses and herpesviruses). Of these 150 MCP sequences, 85 showed higher structural similarities to mirusviruses than to other viruses (Supplementary Fig. S1). Therefore, we considered these 85 MCP sequences as the most appropriate set of sequences for the subsequent phylogenetic analysis. These 85 MCP sequences were derived from 15 assemblies (14 species), including organisms from Alveolata, Amoebozoa, Chlorophyta, Cryptophyta, Rhizaria, and Stramenopiles (Table 1). Most of the homologs showed a bit-score of > 100 and a TM-score to the reference mirusvirus MCPs of > 0.7 (Supplementary Fig. S2). However, homologs from Rhizaria generally displayed low bit-scores, but a wide range of TM-scores to the reference mirusviral MCPs.

**Table 1:** Species and assemblies containing the 85 MCP homologs.

| Species | Major lineage | Assembly | Location of isolation | Environment | Clade |
| --- | --- | --- | --- | --- | --- |
| *Cymbomonas tetramitiformis* | Chlorophyta | GCA_001247695.1 | English Channel | Sea | E01 |
| *Aurantiochytrium* sp. | Stramenopiles | GCA_001462505.1 | Madeira | Sea | E02 |
| *Aurantiochytrium* sp. | Stramenopiles | GCA_003116975.1 | Hiroshima | Sea | E02 |
| *Schizochytrium* sp. | Stramenopiles | GCA_004764695.1 | Missing | Sea | E02 |
| *Parietichytrium* sp. | Stramenopiles | GCA_012862575.1 | Okinawa | Sea | E02 |
| *Hondaea fermentalgiana* | Stramenopiles | GCA_014084085.1 | Mayotte | Mangrove | E02 |
| *Aurantiochytrium acetophilum* | Stramenopiles | GCA_004332575.1 | Biscayne Bay | Mangrove | E02 |
| *Ostreobium quekettii* | Chlorophyta | GCA_905146915.1 | Catanduanes Island | Sea | E02 |
| *Euplotes weissei* | Alveolata | GCA_021440005.1 | Qingdao | Sea | E02 |
| Cryptophyta sp. | Cryptophyta | GCA_026770585.1 | Northern Baffin Bay | Sea | E03 |
| Uncultured Cryptomonadales | Cryptophyta | GCA_947538865.1 | Řimov Reservoir | Fresh water | E03 |
| *Guillardia theta* | Cryptophyta | GCA_000315625.1 | Connecticut | Sea | E03 |
| *Acanthamoeba pearcei* | Amoebozoa | GCA_000826505.1 | Missing | missing | E04 |
| *Paulinella micropora* | Rhizaria | GCA_009731375.1 | Ibaraki | Fresh water | E05 |
| *Paulinella micropora* | Rhizaria | GCA_019918135.1 | Chungnam | Fresh water | E05 |

After all filters were applied, we reconstructed a phylogenetic tree comprising these 85 MCP homologs, together with 79 reference mirusviral MCPs derived from marine metagenomes (Fig. 3). The topology of the subtree of the reference mirusviral MCPs (clades M01 to M07) was generally consistent with a previous report.^4^ The MCP homologs detected in eukaryotic assemblies were not grouped within the reference mirusvirus clades. Instead, they formed five distinct clades, E01 to E05, which were distantly related to the reference MCPs. The representative sequences for the individual clades (i.e., the longest homolog from each clade) displayed predicted 3D structures similar to those of the reference MCP, including the floor domain (Fig. 3). Notably, clades E01–E03 had an additional conserved antiparallel β-strand adjacent to the E-Loop.

**Fig. 3.**
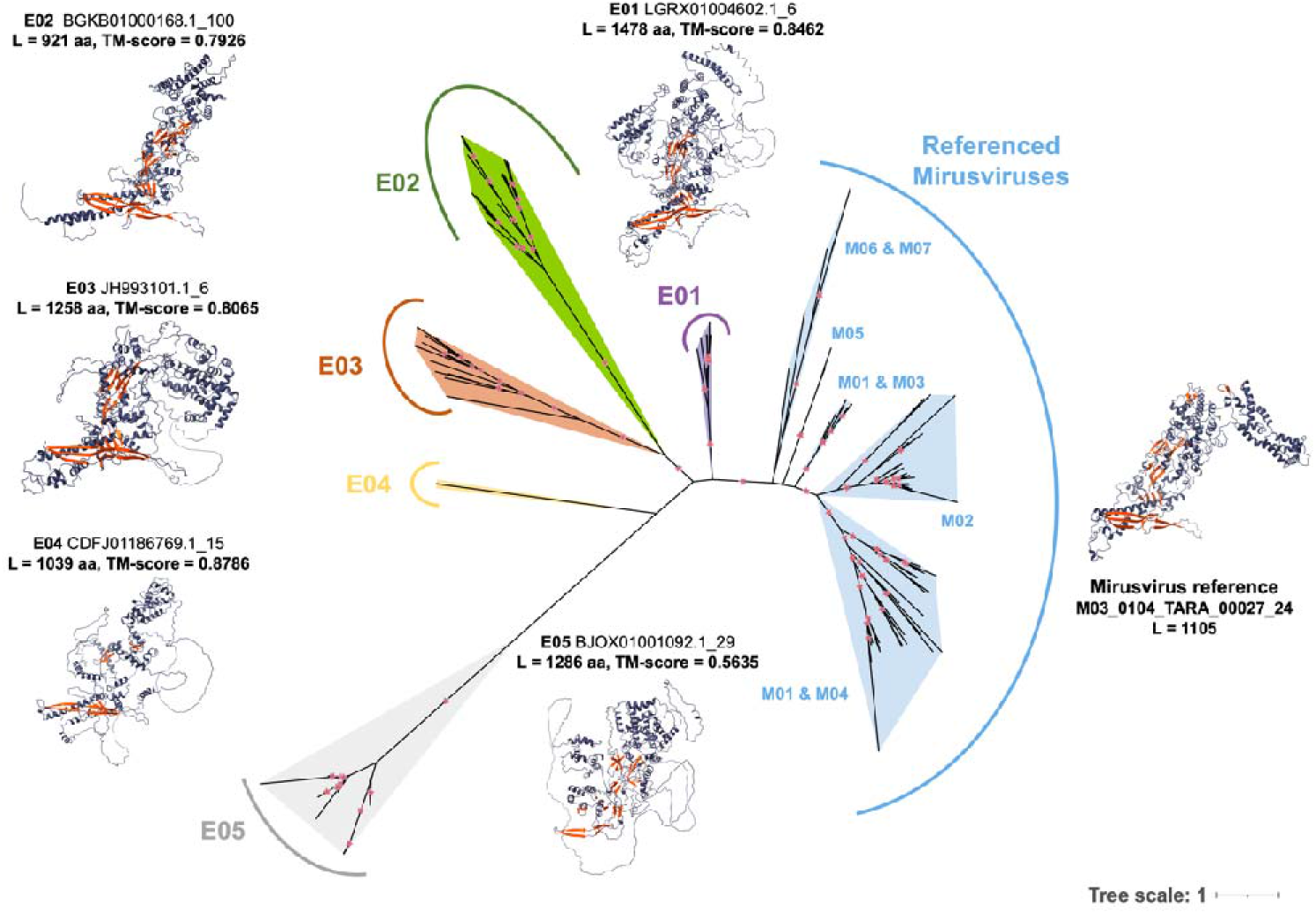
Maximum-likelihood phylogenetic tree of MCPs. Different clades are indicated with different colors: blue represents the reference mirusviral MCPs. Predicted protein structures of the longest homolog within each clade are displayed around the tree. β-sheets are colored orange. Above the structure, the clade number is followed by the scaffold from which the homolog originated and the serial ORF number predicted with prodigal. L, length of this homolog. TM-score is the highest TM-score to mirusvirus reference structures. Small red stars indicate branches with bootstrap support of > 95%. Best-fit model of this tree was Q.pfam+F+R6.

The newly identified clades corresponded to distinct eukaryotic lineages, except Clade E02. Clade E01 consisted of nine homologs from a single genomic assembly of the green alga *C. tetramitiformis*.^11^ Clade E02 was the only clade consisting of homologs (n = 13) from different organisms, including *Euplotes weissei* (Alveolata), *Ostreobium quekettii* (Chlorophyta), and six thraustochytrids (Stramenopiles).^10^ Clade E03 consisted of 18 homologs from three Cryptophyta assemblies. Clade E04 contained a single homolog from an assembly of *Acanthamoeba pearcei* (Amoebozoa). Clade E05 consisted of 44 homologs from two different strains of *Paulinella micropora* (Rhizaria).

### Existence of viral regions and three circular viral genomes

We also investigated the genomic context around the 85 highly conserved MCP homologs. These homologs were found in 83 contigs or scaffolds (hereafter referred to as ‘scaffolds’ for simplicity). The lengths of these scaffolds ranged from 1.2 kilobases to 19.7 megabases. We analyzed all the predicted ORFs on these scaffolds and found that most of the scaffolds encoded many homologs of mirusviral genes, except the very short scaffolds (Fig. 4). Many of the scaffolds showed ORFs showing similarity to eukaryotic homologs. Additionally, most of the scaffolds displayed a higher predicted ORF density than the average level within the same assembly (Supplementary Fig. S3).

**Fig. 4.**
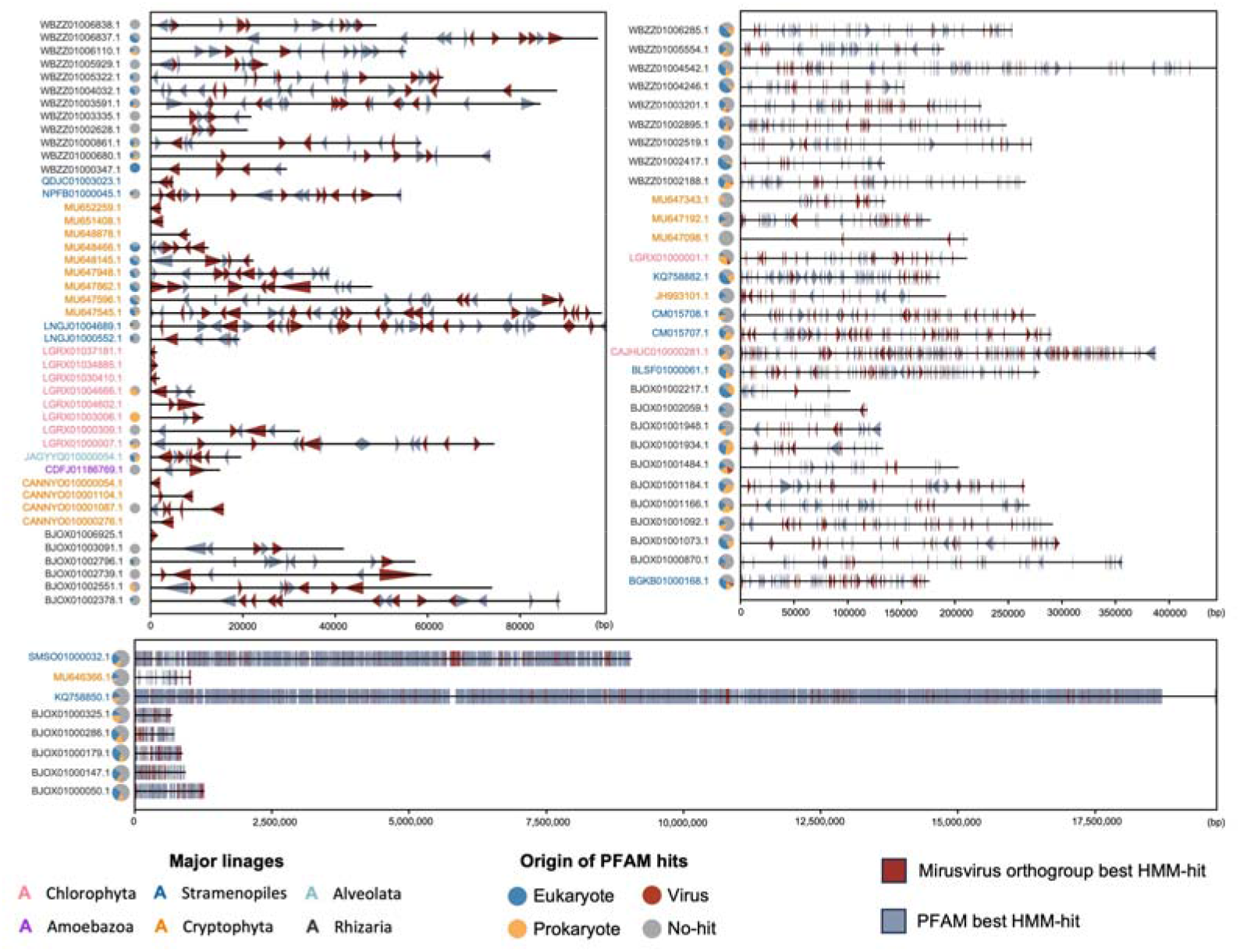
Genomic maps of the 83 scaffolds. Scaffolds were divided into three groups according to their lengths. The color of the text indicates the taxonomic affiliation of each scaffold. The predicted ORFs with mirusvirus orthogroup best hits are colored red. The predicted ORFs with PFAM best hits are colored blue. The tip of the triangle represents the direction of the ORF. Small pie charts represent the proportions of the best-matched organisms obtained by BLAST comparison against the RefSeq database for the ORFs with the PFAM-hits.

From database records at the National Center for Biotechnology Information (NCBI), we found that CM015707.1 and CM015708.1 are circular contigs from *Schizochytrium* sp. (Stramenopiles), with lengths of 289 kilobases and 275 kilobases, respectively. When we examined the remaining 81 scaffolds, we found that CAJHUC010000281.1 (originally 387 kilobases; from *O. quekettii*, Chlorophyta) is also likely to be a circular contig (325 kilobases after circularization) (Supplementary Fig. S4). These circular contigs encode 3–5 mirusvirus markers (of the five selected for this study) and additional functionally important proteins, such as terminases, DNA/RNA polymerases, and heliorhodopsins (Fig. 5). Notably, the three circular genomes lack RNA polymerase subunit B, even though RNA polymerase subunit B is the most commonly detected gene and is conserved in 89% of the reported mirusviral genomes. In contrast, the circular genomes encoded two copies of RNA polymerase subunit A. These circular contigs belonged to Clade E02. A family B DNA polymerase (PolB)-based phylogenetic analysis confirmed the close relationships between these three circular mirusviral genomes (Supplementary Fig. S5). No split genes were detected in these circular contigs, indicating that most genes were intact and not pseudogenized.

**Fig. 5.**
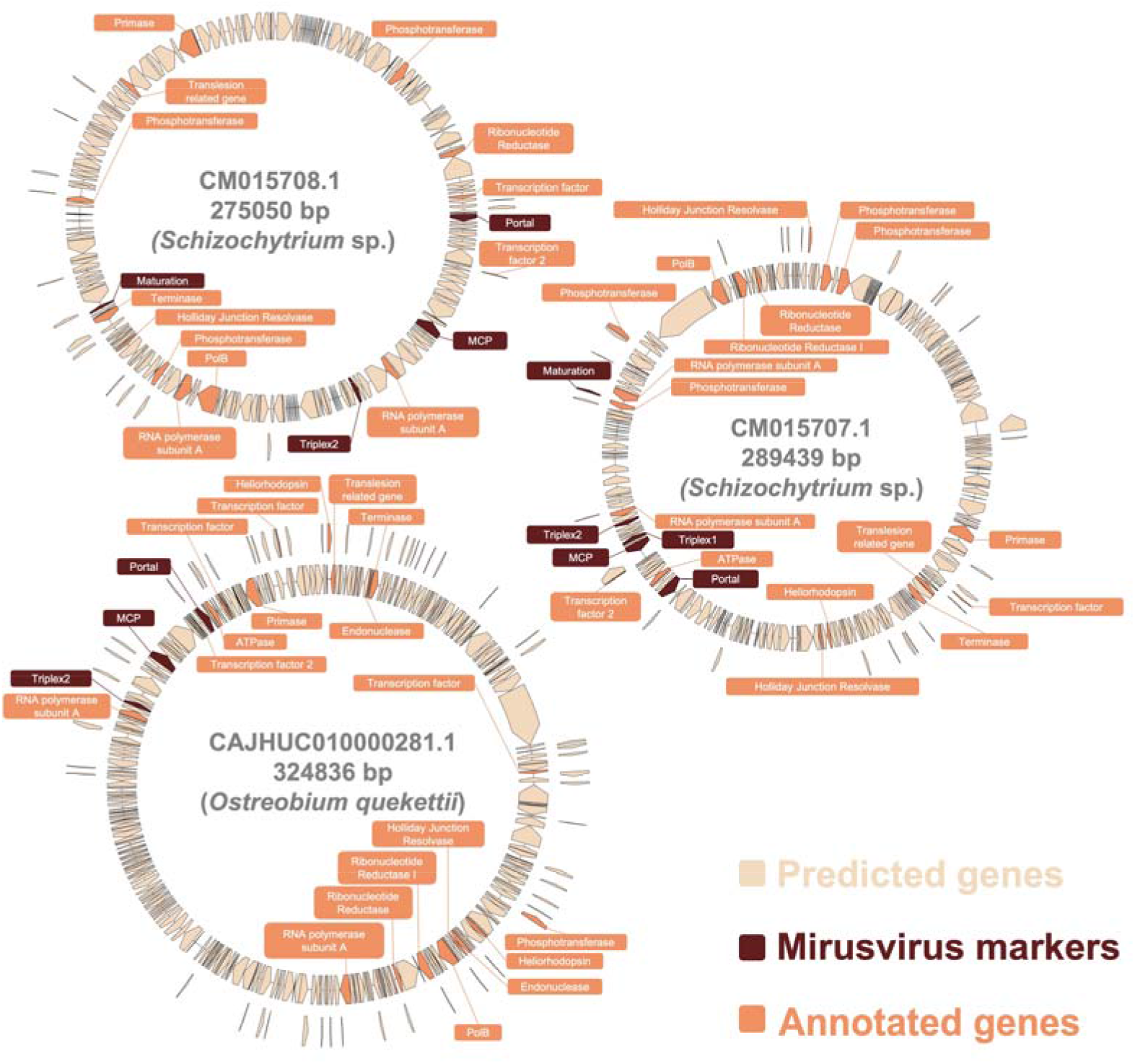
Three circular mirusviral genomes recovered from eukaryotic assemblies. The five markers used in this study are colored brown. Functionally annotated genes are colored orange. Other predicted genes are colored yellowish.

## Discussion

Mirusviruses are reported to be a deep-branching diverse group of dsDNA viruses, but previous studies have provided only limited information on their potential hosts in marine environemnts.^4,10,11^ In the present study, we detected mirusviral maker signals in a large number of eukaryotic (mostly unicellular) genome assemblies (1,348 assemblies; 71% of those analyzed). These eukaryotes included nearly 90% of the eukaryotic genera analyzed and 15 of the 16 major lineages analyzed (Data S2). These potential mirusvirus hosts included not only organisms living in marine environments (e.g., *Aurantiochytrium*, *Euplotes*) but also those living in other environments (e.g., freshwater: *Paulinella, Yamagishiella*; soil: *Dictyostelium*, *Physarum*; parasites of animals or plants: *Trypanosoma*, *Phytophthora*). Therefore, mirusviruses probably infect a broad spectrum of eukaryotes in various types of habitat.

Many of the mirusviral marker sequences identified in eukaryotic assemblies showed low sequence similarity scores to the reference mirusviral sequences (Fig. 2A). This sequence divergence from the reference sequences is probably attributable to the accumulation of mutations (e.g., pseudogenization^19^) after their ancient insertion into the host genomes. This is supported by the numerous abnormally short ORFs for the detected mirusviral markers (Fig. 2B). Despite the presence of decaying genes, 90 genera displayed a comprehensive set of mirusviral marker genes (i.e., four or five markers). Such a high level of marker gene conservation may reflect the relatively recent insertion of the mirusviral genomes into the eukaryotic genomes (Fig. 1).

Our analysis also revealed a wide range of sequence similarity scores, even for ORFs in the normal size range (Fig. 2C). This suggests that factors beyond pseudogenization have contributed to the low sequence similarities between the newly detected sequences and the reference sequences. We analyzed the phylogeny of 85 selected MCP sequences that showed strong similarities (both in sequence and structure) to reference mirusviral MCPs and were thus considered to represent relatively recent insertions (Supplementary Fig. S2). These homologs were only distantly related to the reference mirusviral MCPs and formed five distinct clades (Fig. 3). Therefore, these MCP marker sequences probably originated from viral lineages that are distinct from the previously described mirusviral lineages.

Current research into mirusviruses is based on genomic data from environmental samples or eukaryotes, because no mirusvirus has yet been isolated. Despite their widespread presence in marine environments, the uncertainty regarding their original hosts poses challenges for isolation studies. In this study, we identified culturable eukaryotes showing strong and fresh mirusvirus signals. Notably, a three-stranded antiparallel β-sheet was observed in clade E01, E02, and E03 MCP homologs (Fig. 3). This structure consists of two β-strands in the E-loop at the N-terminus and an additional β-strand at the C-terminus,^1^ and its presence indicates the high structural integrity of the MCP. A previous study reported a circular mirusviral genome in the thraustochytrid *A. limacinum*.^10^ We detected three circular mirusviral genomes, without evidence of pseudogenization, in another thraustochytrid (*Schizochytrium* sp.) and the siphonous green alga *O. quekettii* (Fig. 5). *Schizochytrium* sp. is a marine heterotrophic protist that has been used for commercial production of docosahexaenoic acid-rich oil.^20^ *O. quekettii* is a model organism to study photosynthesis under low light conditions and has an endolithic lifestyle in association with all sorts of marine limestones such as coral skeletons.^21^ The circular status of the genomes suggests that these mirusviruses either latently^10,22^ or persistently^23^ infect these hosts. Eukaryotes harboring circular mirusviral genomes or high-integrity MCP sequences (E01–E03 in Table 1) represent promising candidates for the isolation of mirusviruses because their infections are probably recent or on-going events.

Viral genes are known to contribute to the critical evolution of their hosts.^24,25^ The invasion of eukaryotic host genomes by DNA viral genomes is widespread among unicellular eukaryotes.^13^ In particular, large dsDNA viruses have been reported to form giant viral endogenous elements within their host genomes.^26,27^ The detection of mirusvirus signals in various eukaryotes indicates that mirusviruses are not an exception to this phenomenon (Data S1, S2). We identified multiple mirusvirus homologs in the scaffolds that harbored mirusviral MCPs, and many eukaryotic genes were also discovered in these scaffolds (Fig. 4). These suggest that the genomic regions are giant endogenous viral elements originating from mirusvirus genomes. It has been suggested that giant viral endogenous elements confer unique capabilities upon their hosts (usually unicellular eukaryotes) by providing various genes of viral origin,^26^ in a similar way to endogenous viral elements in higher eukaryotes.^28,29^ Given the extensive presence of mirusvirus-like genomic regions in eukaryotic genomes, mirusviruses may have contributed to the genomic innovation of their hosts.^30^

## Methods

### Unicellular eukaryotic genome assembly data

We compiled the unicellular eukaryotic genome assembly data by retrieving all the relevant eukaryotic genome assemblies from the GenBank database (as of June 2023). Assemblies from Fungi (NCBI Taxonomy ID 4751), Metazoa (NCBI Taxonomy ID 33208), and Streptophyta (NCBI Taxonomy ID 35493) were excluded. A total of 1,901 assemblies were collected for subsequent analysis. Organisms that were not classified at the genus level were treated as distinct genera. To identify mirusvirus-originating sequences within these assemblies, gene calling was performed on all compiled data with the Prodigal v2.6.3 software, with the parameter ‘-p meta’.^31^ Predicted sequences shorter than 20 amino acids or longer than 4,000 amino acids were discarded.

### Creation of mirusvirus marker protein models

The reference metagenome-assembled mirusvirus genomes were obtained from a previous study.^4^ Gene calling was performed with Prodigal, and orthologous groups were then generated with Orthofinder v2.5.2.^32^ The core gene orthologous groups were annotated with a BLASTp search against the RefSeq database with Diamond v2.1.8,^33^ and subsequent manual curation. For the five virion module protein markers, redundant sequences with > 90% sequence identity were removed from the five orthologous groups with cd-hit^1^ v4.8.1 and the parameter ‘-c 0.9’.^34^ The HMMs of the five markers were built with HMMER v3.3.2. ^35^

### Detection mirusvirus marker sequences in eukaryotic assemblies

To detect mirusviral marker sequences within the eukaryotic assemblies, we used HMMs corresponding to five marker genes. We screened our predicted protein database, retaining hits with an E-value < 10^−5^ as determined with HMMER. To ensure the specificity of our results for mirusviruses, we established additional controls to exclude similar sequences from other viruses. First, we downloaded caudovirus and herpesvirus protein sequences from NCBI Virus (as of October 26, 2023). We then curated a dataset of homologs for the selected marker genes (for caudoviruses: MCP, Portal, Maturation; for herpesviruses: MCP, Triplex1, Triplex2, Portal, Maturation) using keyword filtering based on the NCBI annotations. Sequence redundancy was removed with cd-hit, with a cut-off of 90% sequence identity. We then constructed HMMs for each of the caudovirus and herpesvirus proteins and used these models to examine our mirusvirus hits. Any protein sequences from the eukaryotic assemblies that demonstrated a lower E-value when matched with the caudovirus or herpesvirus protein models than when compared with the mirusvirus models were excluded from further analysis. We also used hmmscan to scan the remaining mirusvirus hits against HMMs in the PFAM database (as of November 7, 2023),^36^ and discarded hits with lower E-values against any PFAM HMM.

### 3D structural analyses of MCP homologs

Protein 3D structural predictions were made with AlphaFold v2.3.2 (-t 2023-11-14).^37^ For structural comparisons with known HK97-fold MCPs, we also predicted the 3D structures of reference MCP sequences from mirusviruses, herpesviruses, and caudoviruses. Sequences shorter than 200 amino acids or with > 70% sequence identity to other sequences in the same viral group were removed from the reference MCP sequences. In total, 37 herpesvirus MCP models and 252 caudovirus MCP models were referenced. Structural similarities between proteins were calculated with Foldseek v6-29e2557 with the program ‘easy-search’.^38^ Protein structures were visualized with ChimeraX v1.7.^39^

### Phylogenetic analysis

Multiple-sequence alignments of MCP and PolB sequences longer than 200 amino acids were generated with Clustal-Omega v1.2.4.^40^ We trimmed the alignments with trimAl v1.4.1, with the parameter ‘-gt 0.1’.^41^ Maximum-likelihood phylogenetic trees were constructed with IQ-TREE v2.2.2.6,^42^ with 1,000 ultrafast bootstrap replicates. Models were selected with ModelFinder.^43^ The phylogenetic trees were visualized with iTOL v6.8.1.^44^ For the phylogenetic tree of PolB, we collected all mirusviral PolB homologous sequences on those 83 scaffolds and also used eukaryotic reference sequences from a previous study.^45^

### Annotation of contigs and scaffolds

We identified mirusvirus orthologous groups that were shared by more than 10 reference mirusviral metagenome-assembled genomes. Using the HMMs of these orthologous groups as the queries, we scanned all 83 scaffolds (E-value < 10^−3^). We also used PFAM models as queries to scan these scaffolds (E-value < 10^−3^). We annotated the related organism for those ORFs with a best hit in PFAM by using Diamond BLASTp against the RefSeq database (E-value < 10^−3^). For the predicted genes that showed similarities to both mirusvirus orthologous groups and PFAM models, the annotation result was based on the hit with the lower E-value. To determine whether these scaffolds were likely to be circular, we performed a similarity comparison of different parts of the same sequence with NCBI web-based versions of BLASTn and DigAlign to determine whether the extremities of the sequence showed the same sequence (https://www.genome.jp/digalign/).^46^ For the three circular mirusvirus contigs, we re-predicted their ORFs with the default parameters of Prodigal. The genomic maps of the circular contigs were generated with the Python Package ‘DNA features viewer’,^47^ and the annotations were based on the best hits of the predicted ORFs to the mirusvirus orthogroups (E-value < 10^−3^). To investigate pseudogenization, we used Diamond BLASTp to query all ORFs derived from the circular genomes against the RefSeq database, using the parameter ‘-k 3 -e 10’. We looked for adjacent ORFs showing similarity to different parts of the same reference sequence as potential signals of pseudogenization.

## Acknowledgments

We thank Junyi Wu and Ruixuan Zhang for valuable discussions. This work was supported, in part, by JSPS/KAKENHI (nos 22H00384 and 19H05667). Computational time was provided by the Supercomputer System, Institute for Chemical Research, Kyoto University (Kyoto, Japan). Molecular graphics and analyses were performed with UCSF ChimeraX, developed by the Resource for Biocomputing, Visualization, and Informatics at the University of California, San Francisco (CA, USA), with support from National Institutes of Health (R01-GM129325) and the Office of Cyber Infrastructure and Computational Biology, National Institute of Allergy and Infectious Diseases (MD, USA). We thank Edanz (https://jp.edanz.com/ac) for editing a draft of this manuscript.

## Author Contributions

HZ designed the work, performed all the analyses, and wrote the initial draft of the manuscript. LM contributed to the data preparation and assisted with the analyses. HH and HO supervised HZ and contributed to the finalization of the manuscript. All authors have read and approved the final version of the manuscript.

## Data and Code Availability

- The annotations, predicted ORFs, HMM files, alignments, predicted structure files, and tree files are available through GenomeNet (https://www.genome.jp/ftp/db/community/Mirusvirus_host_range/).
- This paper does not report original code.
- Any additional information required to reanalyze the data reported in this paper is available from the authors upon request.

