## Supplementary for "Eukaryotic genomic data uncover an extensive host range of mirusviruses"

**Data S1: Accession numbers and marker distribution of the 1901 used genomic assemblies, related to Figure 1, Table S1 and Methods.**

**Data S2: Taxonomic assignments of the eukaryotic assemblies containing detected mirusvirus markers related to Figure 1, Table S1.**

**Supplementary Table S1: Information of collected 1901 eukaryotic genomic assemblies.**

| Major lineage | Number of genera | Number of assemblies |
|---|---|---|
| Stramenopiles | 83 | 668 |
| Alveolata | 73 | 526 |
| Chlorophyta | 44 | 226 |
| Discoba | 28 | 207 |
| Amoebazoa | 22 | 79 |
| Rhizaria | 10 | 61 |
| Metamonada | 12 | 53 |
| Rhodophyta | 17 | 42 |
| Opisthokonta | 12 | 17 |
| Haptista | 6 | 9 |
| Cryptophyta | 7 | 8 |
| Apusozoa | 1 | 1 |
| Breviatea | 1 | 1 |
| Glaucophyta | 1 | 1 |
| Malawimonadida | 1 | 1 |

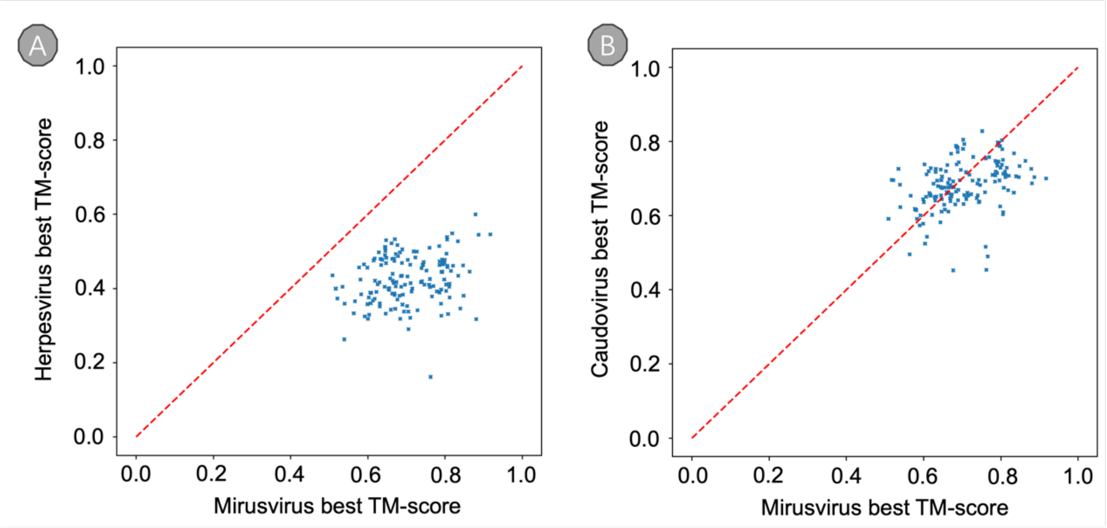


**Supplementary Fig S1: TM-scores of detected 150 MCP sequences for different duplodnaviruses. (A)** Highest TM-score of the MCP homologs compare with mirusviruses and herpesviruses, respectively. **(B)** Highest TM-score of the MCP homologs compare with mirusviruses and caudoviruses, respectively. **(A-B)** Each dot represents a MCP homolog. The slope of the red dashed line is 1.

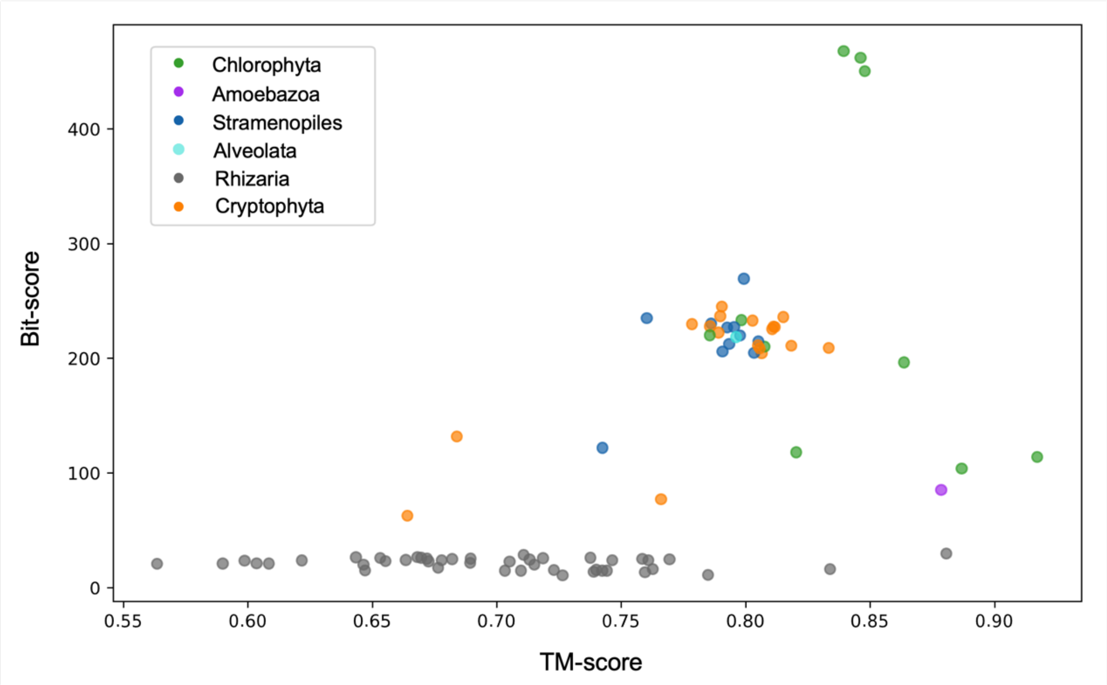


**Supplementary Fig S2: Relationship between bit-scores and TM-scores of the 85 MCP homologs.** Each dot represents a MCP homolog. X-axis represents the highest TM-score of homologs compared with all mirusvirus references. Y-axis represents the hmmscan bit-scores against the reference mirusvirus MCP model. Color of each dot represent the taxonomic groups of the genomic assemblies.

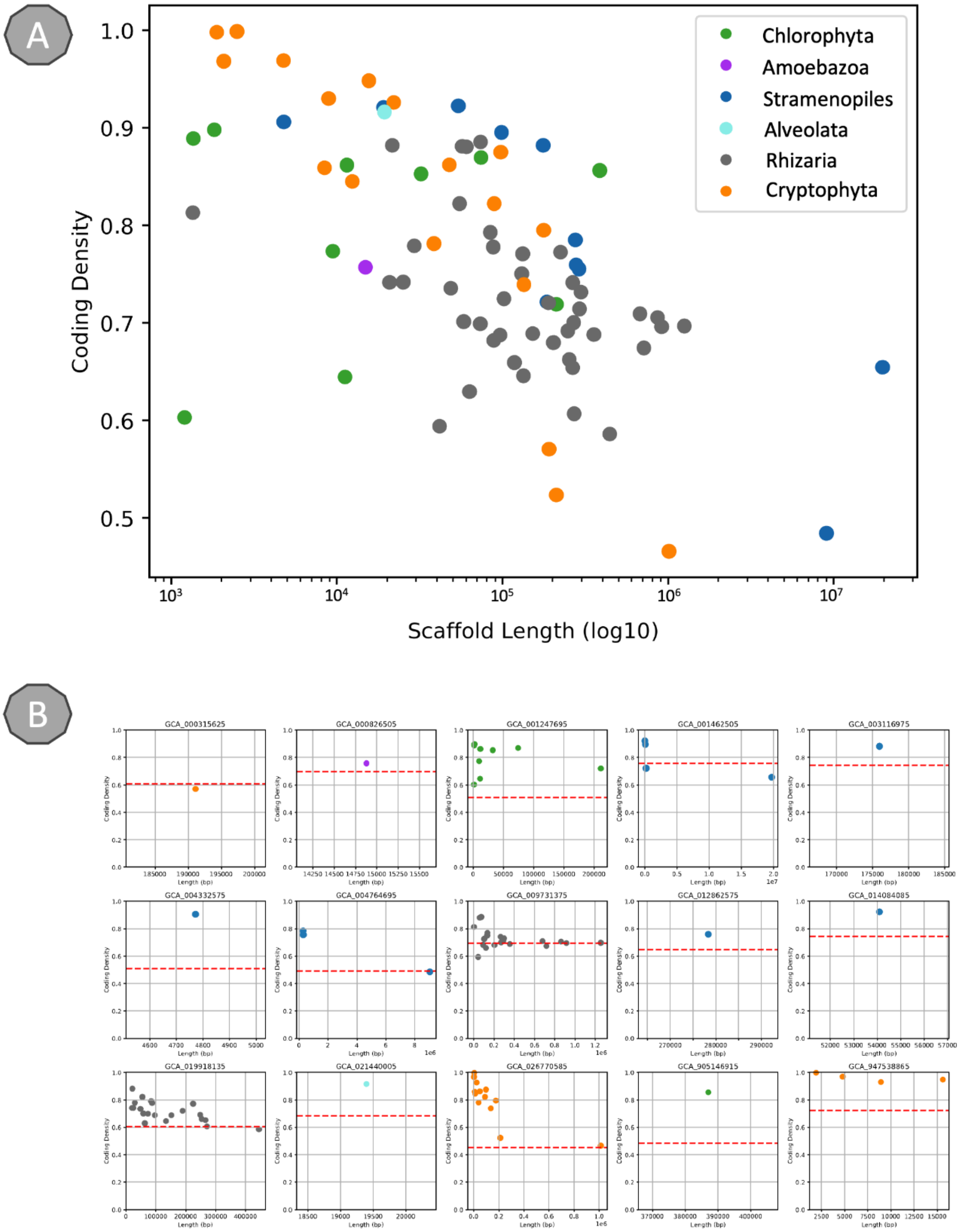


**Supplementary Fig S3: Predicted ORF density of these 83 contig and scaffolds. (A)** Each dot represents a contig or scaffold. X-axis represents the length of scaffold or contig. Y-axis represents the predicted ORF density of this contig or scaffold. Color of each dot represent the taxonomic origin of the genomic assembly of the identified homolog. **(B)** Each sub-figure represents an assembly, and each dot represents a contig or scaffold. The red dushed line means the average predicted ORF density of this assembly. The color is same with (A).

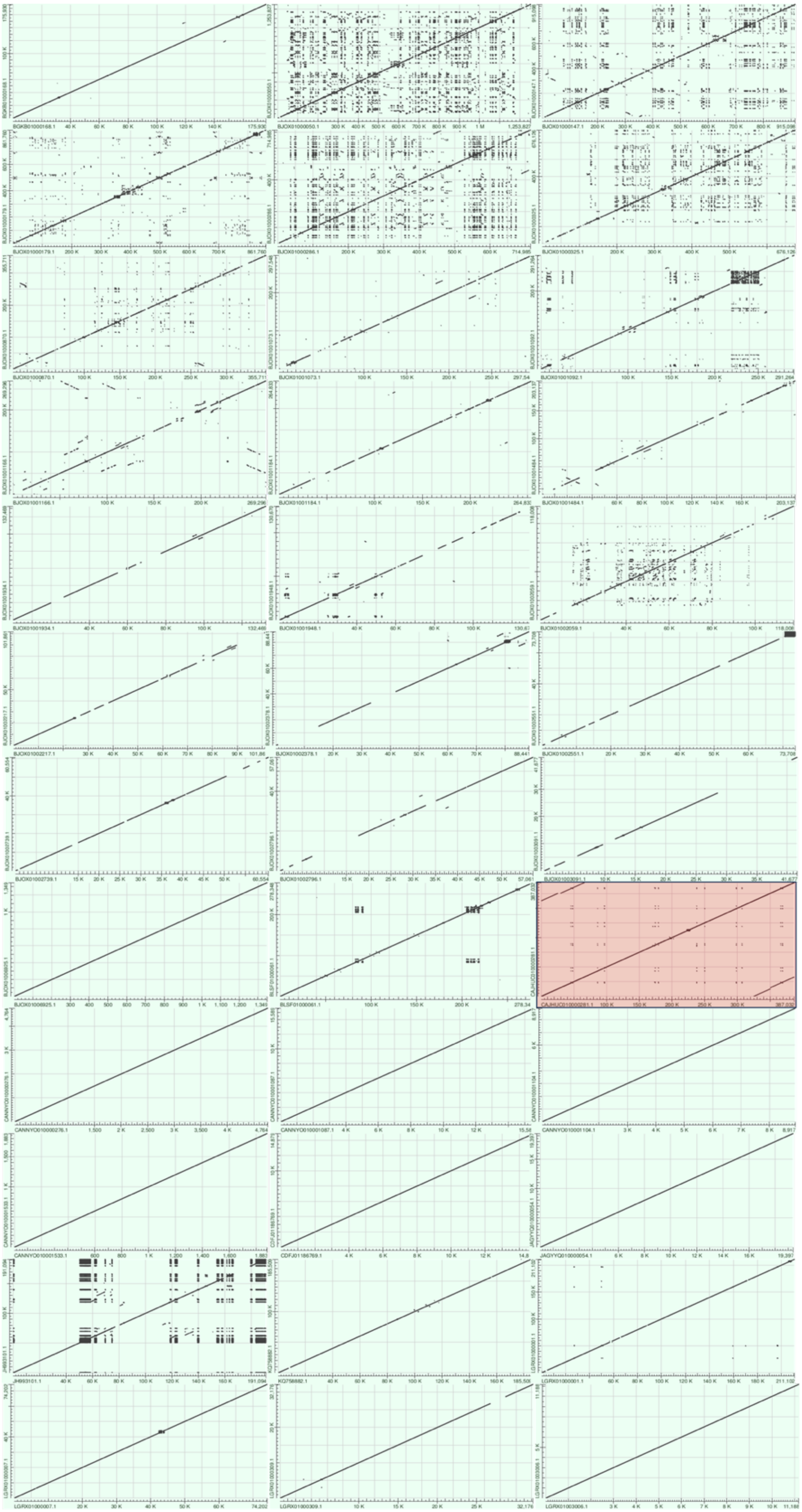

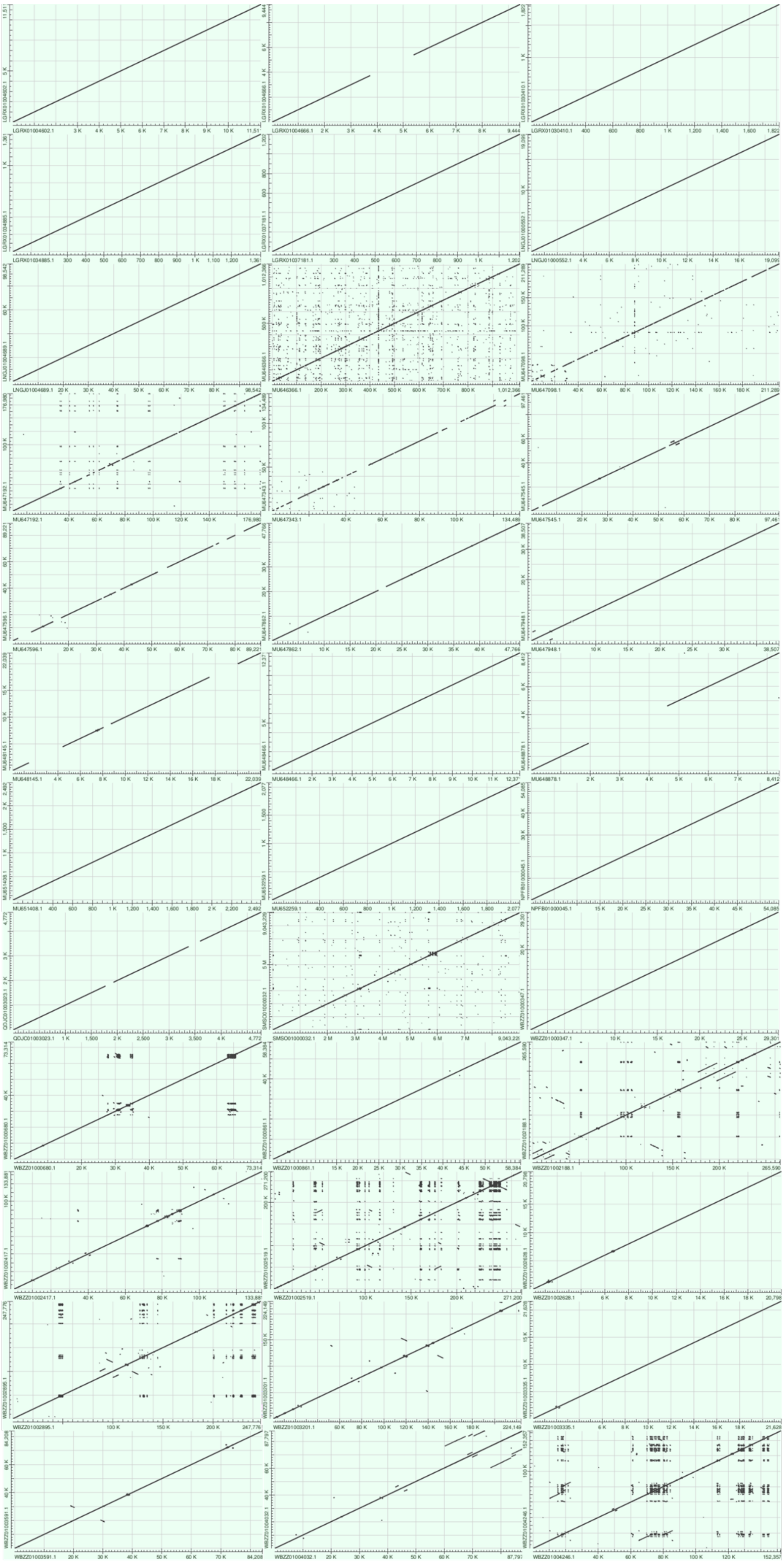
LGRX01004602.1
LGRX01004666.1
LGRX01030410.1
LGRX01034985.1
LGRX01037181.1
LNGJ01000552.1
LNGJ01004689.1
MU646366.1
MU647098.1
MU647192.1
MU647343.1
MU647545.1
MU647596.1
MU647862.1
MU647948.1
MU648145.1
MU648466.1
MU648878.1
MU651408.1
MU652259.1
NPFB01000045.1
QDJC01003023.1
SMSO01000032.1
WBZZ01000347.1
WBZZ01000680.1
WBZZ01000861.1
WBZZ01002188.1
WBZZ01002417.1
WBZZ01002519.1
WBZZ01002628.1
WBZZ01002895.1
WBZZ01003201.1
WBZZ01003335.1
WBZZ01003591.1
WBZZ01004032.1
WBZZ01004246.1

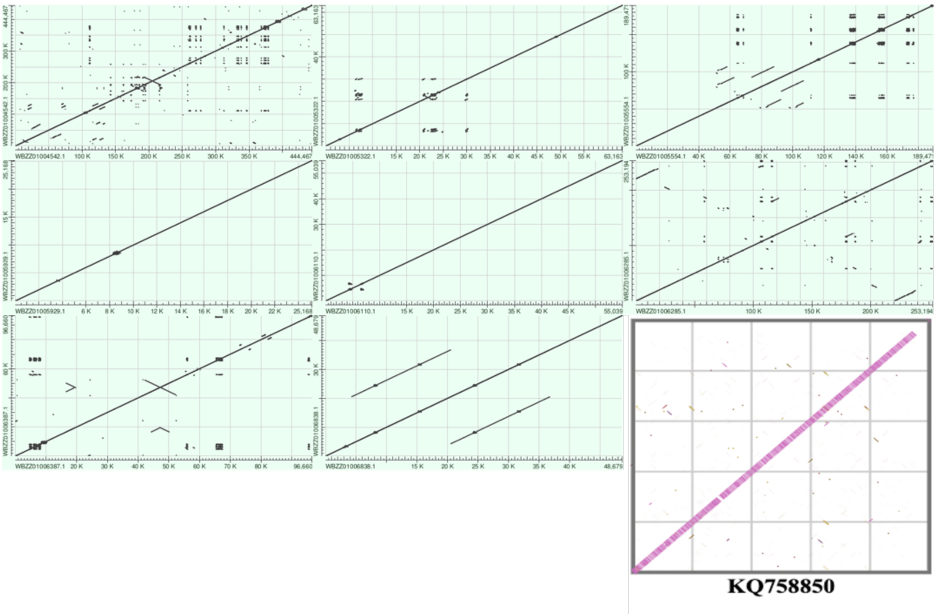


**Supplementary Fig S4: Sequence similarity of the 81 contigs and scaffolds when compared with themselves.** All the sub-figures are the similarity of different location within a scaffold or contig. CAJHUC010000281.1 was marked by red square, since the similarity of its first 60 kilobases and last 60 kilobases. All subfigures were generated by NCBI BLAST website, except the subfigure of KQ758850 was generated by DigAlign due to its length.

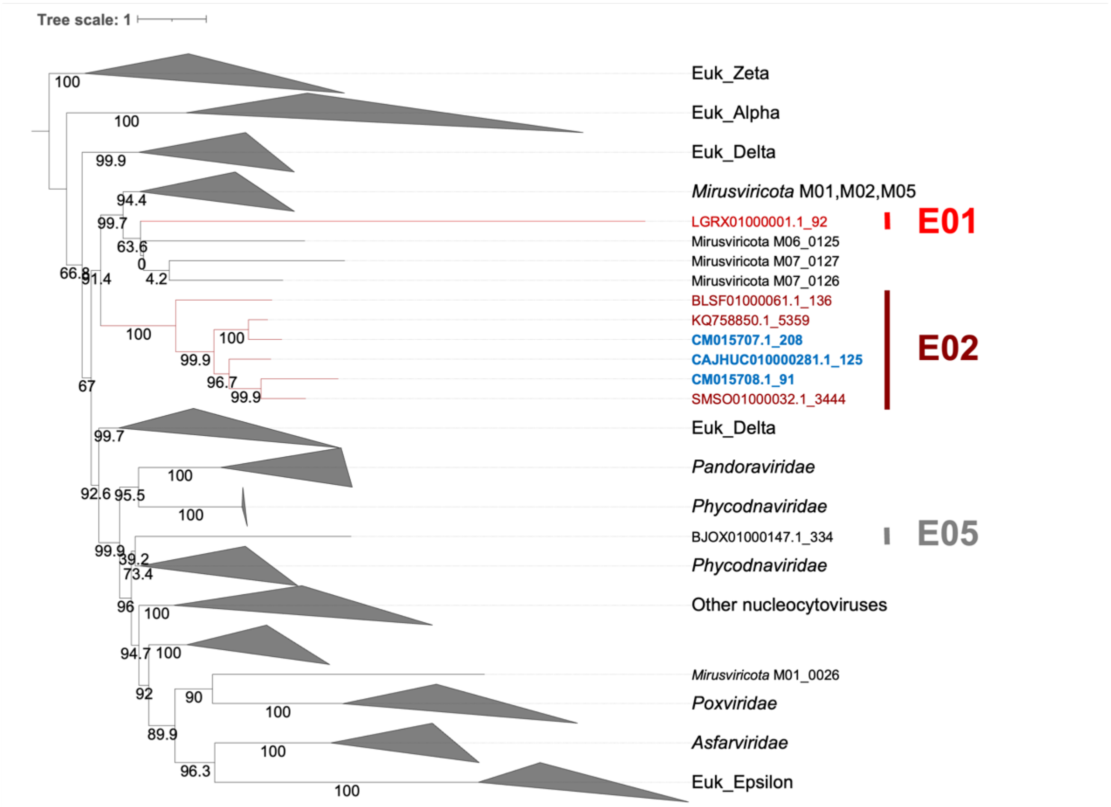


**Supplementary Fig S5: Maximum-likelihood phylogenetic tree of PolB.** *Mirusviricota*-like homologs were colored in red. The root of the tree was chosen at the midpoint. Ultrafast bootstrap support values are provided along the branches. The clade of MCP located on the scaffold was annotated in the right. PolBs from the three circular contigs were colored in blue. The best-fit model of this tree is Q.pfam+F+R10.
